## Supplementary Figures and Tables for "ALS-FUS mutation affects the activities of HuD/ELAVL4 and FMRP leading to axon phenotypes in motoneurons"

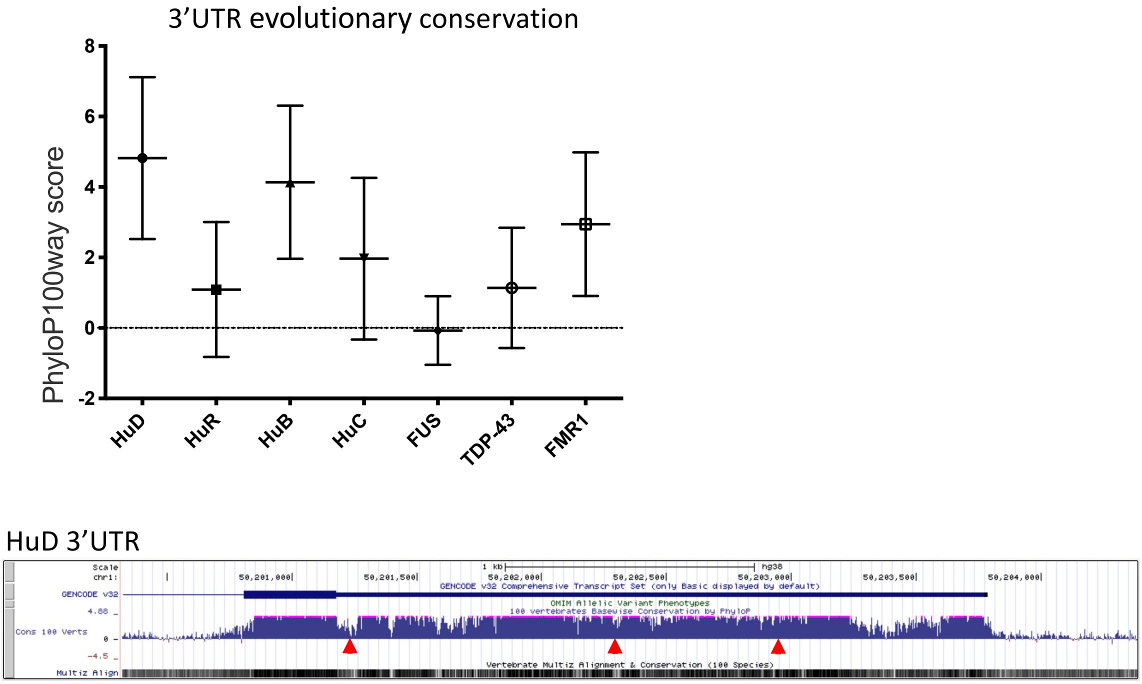


**Supplementary Figure S1. Conservation of HuD/ELAVL4 3'UTR**

Top: graph showing the mean and standard deviation values of the phyloP100way score relating to the 3'UTR sequences of indicated RBP genes (http://genome.ucsc.edu; UCSC Genome Browser assembly ID: hg38). Bottom: screenshot of the UCSC genome browser (http://genome.ucsc.edu; UCSC Genome Browser assembly ID: hg38) showing the sequence conservation of the HuD 3'UTR. Binding sites of miR-375 are indicated by red arrowheads.

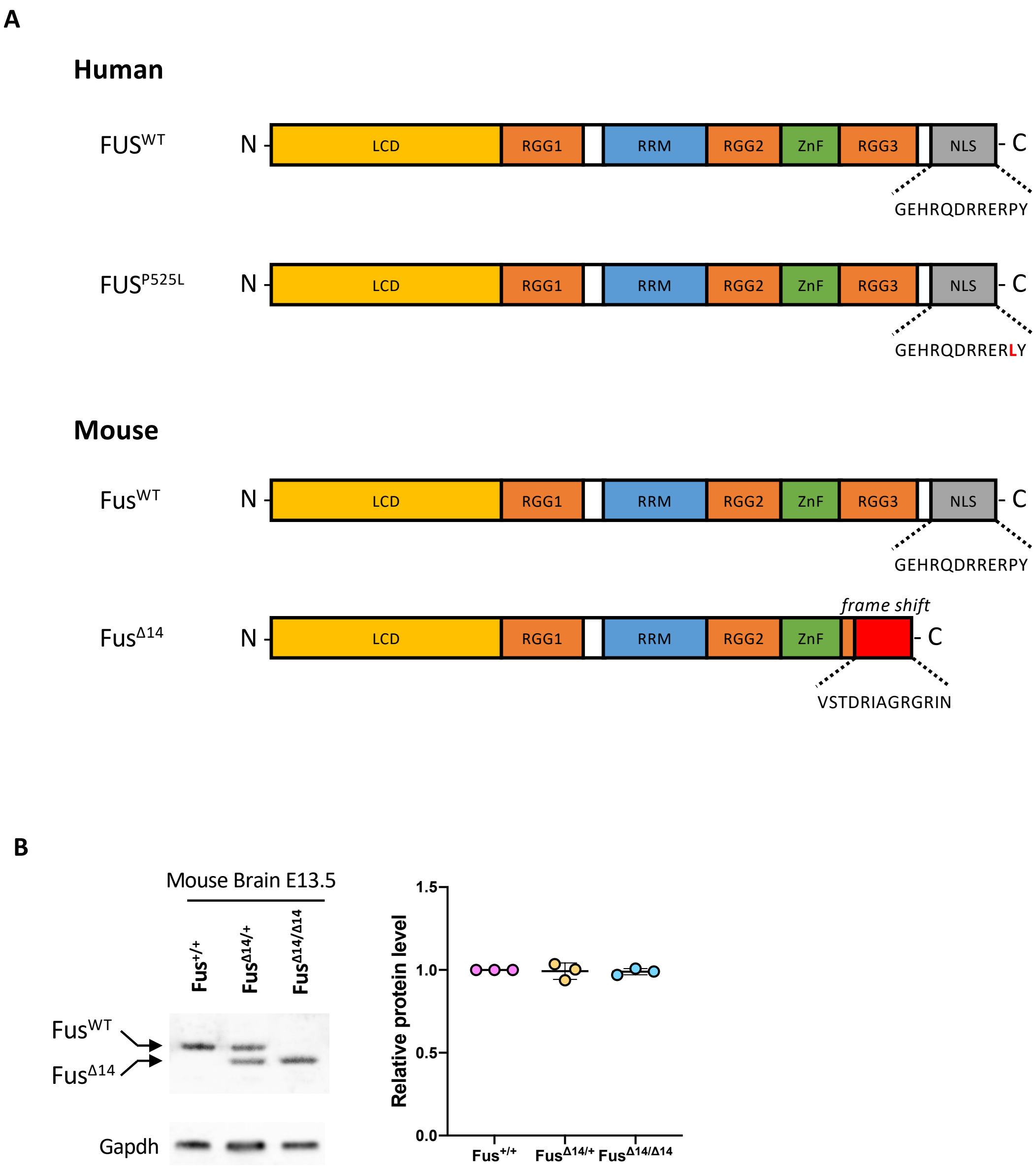


**Supplementary Figure S2. FUS mutants used in this study**

(A) Schematic representation of human and mouse FUS mutants used in this study. (B) Western blot analysis and quantification of wild-type and mutant Fus protein levels in the brain of mouse models used in this study at E13.5.

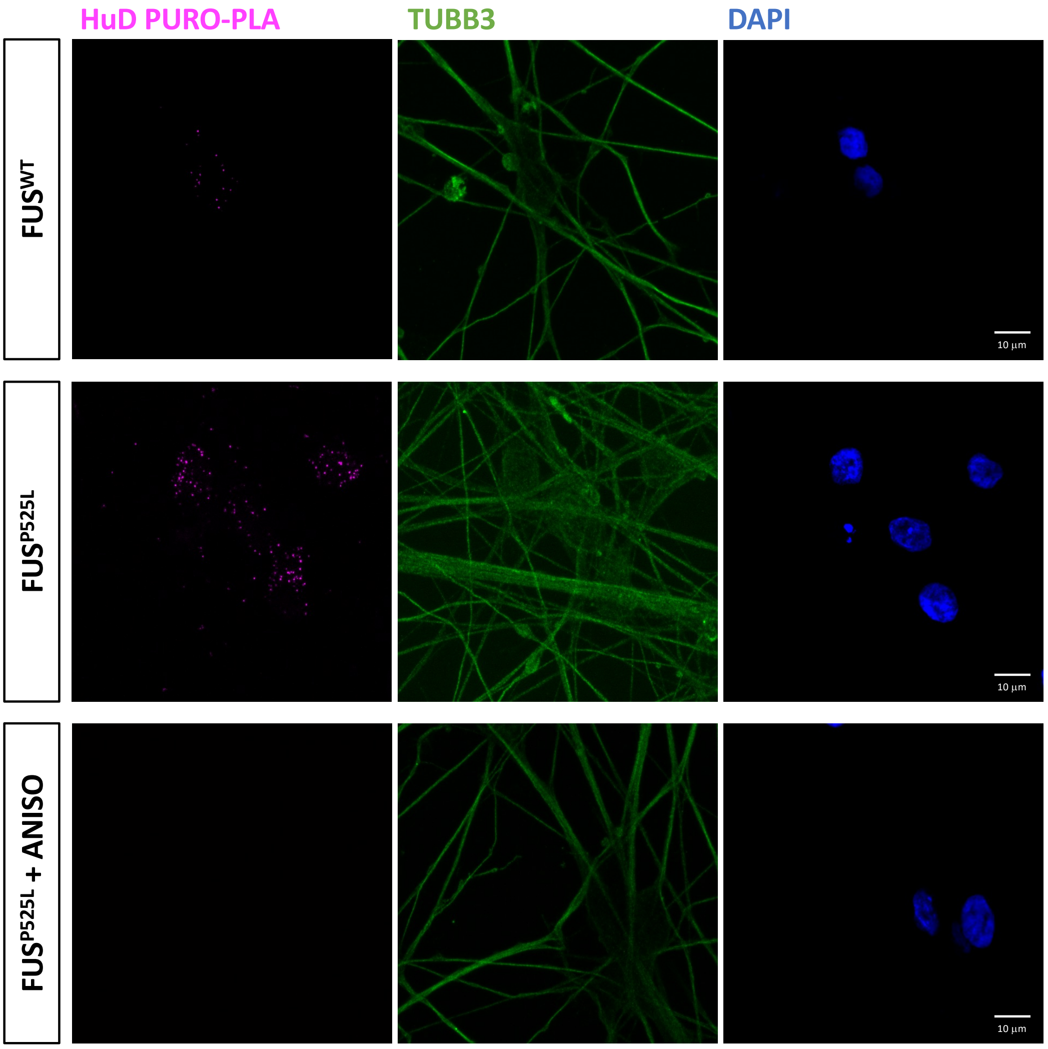


**Supplementary Figure S3. HuD PURO-PLA and TUBB3 immunostaining analysis**

Single panels of the PURO-PLA (HuD, magenta) and immunostaining (TUBB3, green) analysis in FUS$^{WT}$ and FUS$^{P525L}$ hiPSC-derived spinal MNs, shown in Figure 2. DAPI (blue) was used for nuclear staining. Cells treated with the eukaryotic protein synthesis inhibitor anisomycin (FUS$^{P525L}$ + ANISO), used as a PURO-PLA control, are also shown. Scale bar: 10 µm.

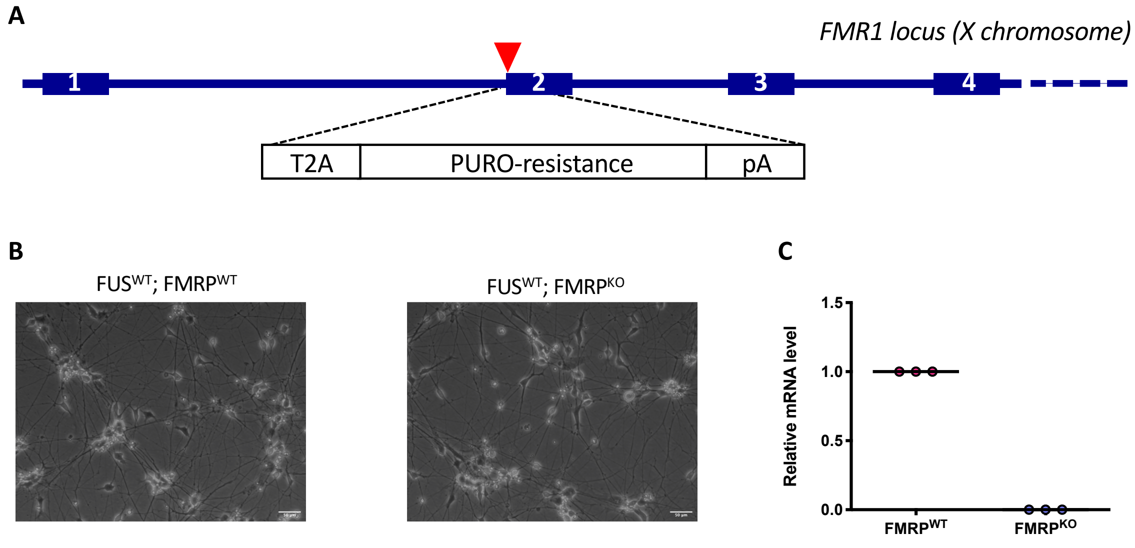


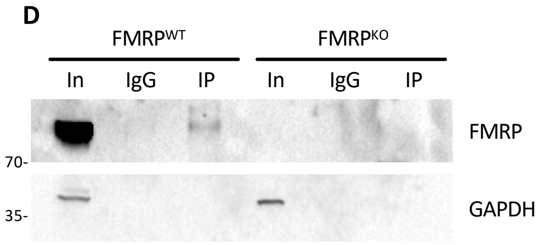


| Target | Percentage of input in FMRP$^{KO}$ RIP |
|---|---|
| *HuD* | 0.0002 |
| *MAP1B* | 0.2580 |
| *ATP5O* | 0.0006 |

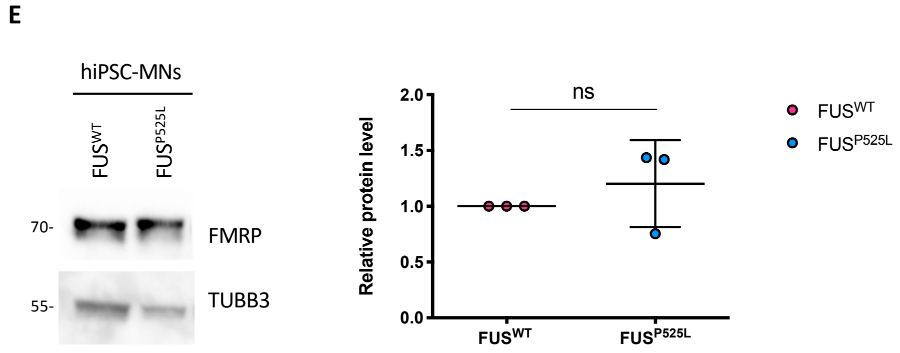


**Supplementary Figure S4. FMR1 knock-out hiPSCs**

(A) Schematic representation of the FMR1 locus and the strategy for CRISPR/Cas9 knockout. The red triangle indicates the target site of the CRISPR guides. The cassette encoding for the self-cleavage peptide T2A, the puromycin resistance gene (PURO-resistance) and the BGH cleavage and polyadenylation site (pA), schematized in the figure, was integrated in the genome upon homology directed repair, interrupting the FMR1 coding sequence. Selection with puromycin was followed by clonal expansion and characterization of the FMR1 knock-out line, as described in [36]. (B) Phase contrast images of hiPSC-derived MNs obtained from the parental and the FMRP knock-out targeted lines. Scale bar: 50 µm. (C) Analysis of the FMR1 mRNA levels by real time qRT-PCR in $FMRP^{WT}$ and $FMRP^{KO}$ hiPSC-derived MNs. (D) Left: western blot analysis of the FMRP RIP assay, as in Figure 2A, performed on hiPSC-derived MNs obtained from the parental and the FMRP knock-out targeted lines. The molecular weight is indicated on the left. Right: table showing the results of the FMRP RIP analysis of MNs derived from the FMR1 knock-out line, used as a negative control of the experiments shown in Figure 2A,B. (E) FMRP protein levels analysis by western blot in $FUS^{WT}$ and $FUS^{P525L}$ hiPSC-derived spinal MNs. The molecular weight is indicated on the left. The graph shows the average from 3 independent differentiation experiments, error bars indicate the standard deviation (Student's t-test, paired, two tails, ns: $p > 0.05$). TUBB3 signal was used for normalization.

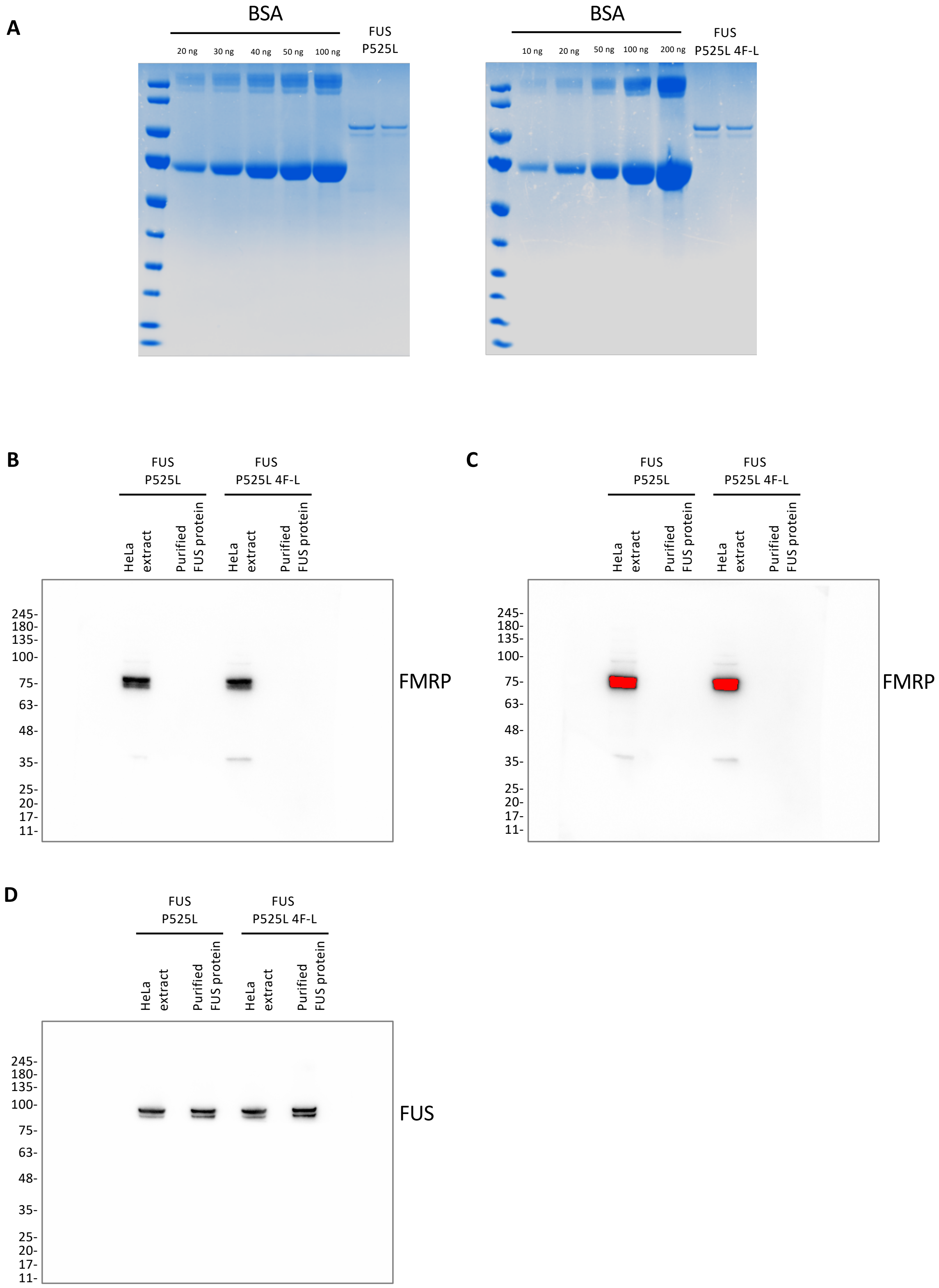


**Supplementary Figure S5. Purification of the recombinant FUS protein from HeLa extracts**

(A) Recombinant flag-tagged FUS proteins (left: P525L; right: P525L 4F-L) were purified from HeLa extracts and run on an electrophoretic gel together BSA at known concentrations for quantification.

Coomassie blue staining of the gels confirmed effective purification of the recombinant proteins. (B-D) Western blot analysis of FMRP (B,C) and recombinant flag-tagged FUS (D; using an anti-flag antibody) in HeLa extracts (8 µg) and purified recombinant flag-tagged FUS protein samples (3 ng). Panel (C) shows the same filter as in (B) at higher exposure time, beyond saturation of the FMRP signal in HeLa extract samples. This analysis confirms no detectable contamination of FMRP in purified recombinant flag-tagged FUS protein samples from HeLa extracts.

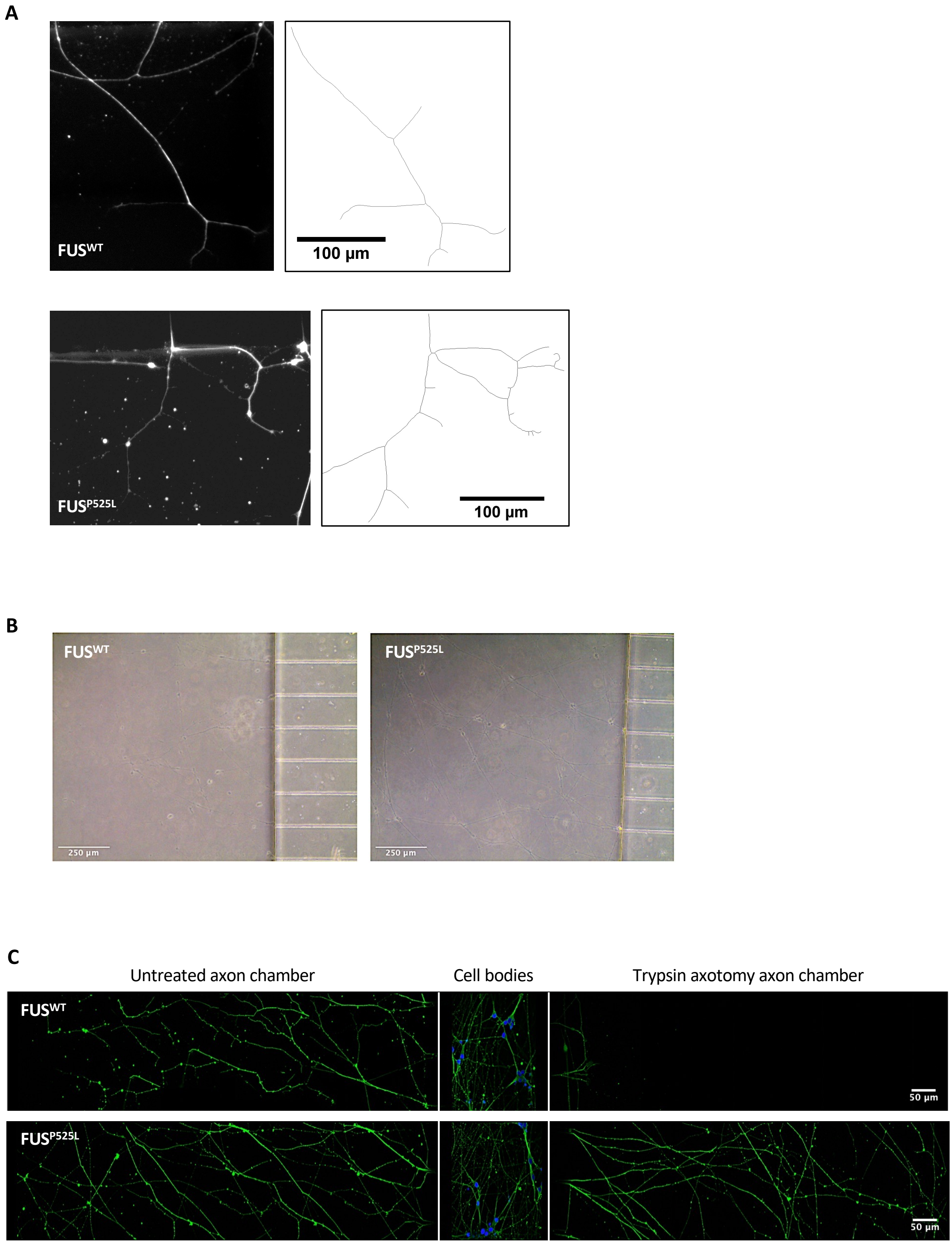


**Supplementary Figure S6. Axon branching analysis and axotomy and recovery assays**

(A) Immunofluorescence images (TUBB3 in white; left) and corresponding images generated with the Skeleton plugin of ImageJ (right), showing axons of $FUS^{WT}$ and $FUS^{P525L}$ human iPSC-derived MNs in the axon chamber of compartmentalized chips. Scale bar: 100 μm. (B) Brightfield images of the axons, before trypsin axotomy, of $FUS^{WT}$ and $FUS^{P525L}$ human iPSC-derived MNs in the axon chamber of compartmentalized chips. Scale bar: 250 µm. (C) Immunostaining of TUBB3 (green) in $FUS^{WT}$ and $FUS^{P525L}$ hiPSC-derived spinal MNs cultured in compartmentalized chips and allowed to recover for 30 hours after the trypsin treatment to induce axotomy. DAPI (blue) was used for nuclear staining. The figure shows the trypsin treated axon chamber on the right, the untreated axon chamber, used as control, on the left and the cell body chamber in the middle. Scale bar: 50 µm.

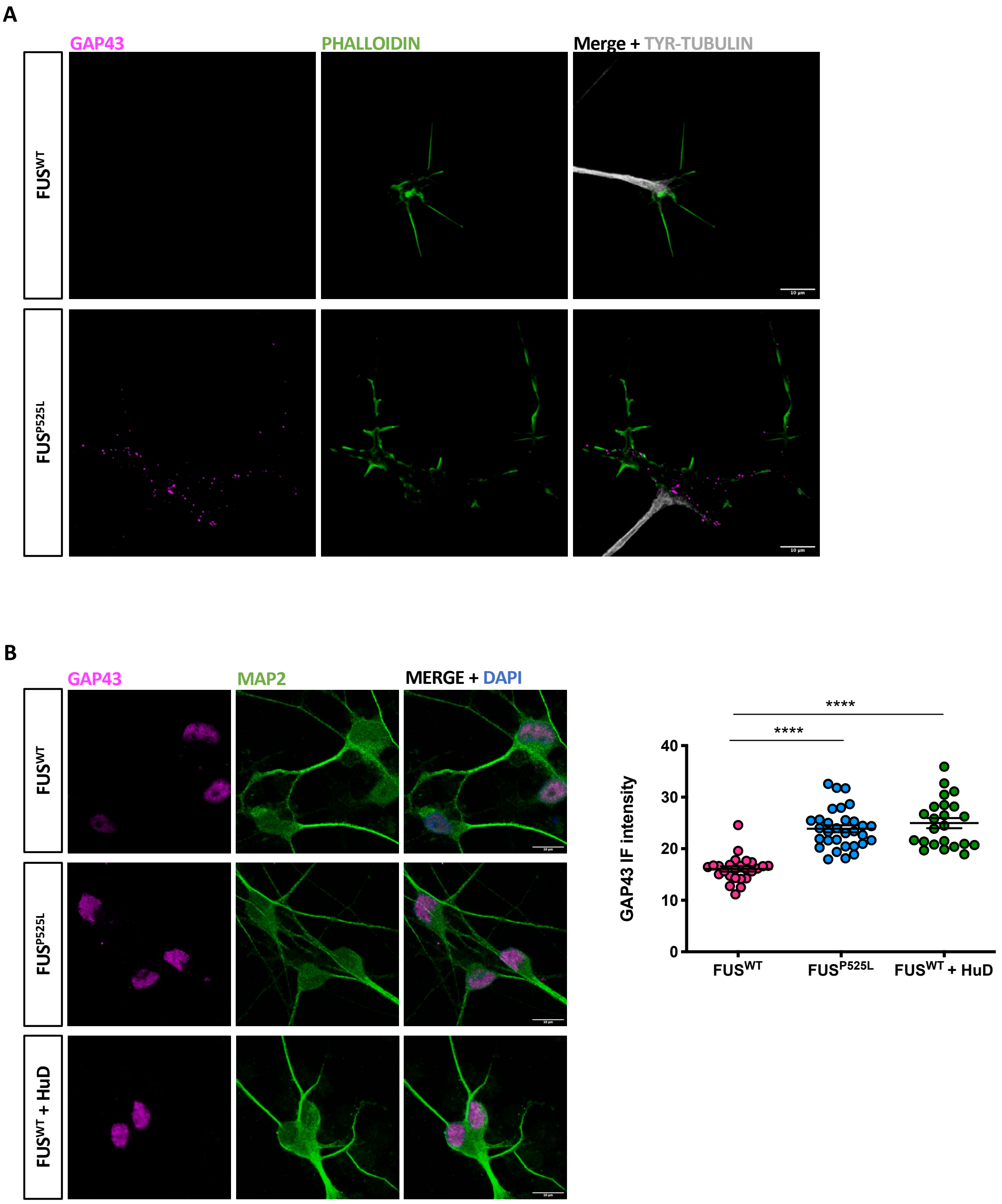


**Supplementary Figure S7. Immunostaining analysis of GAP43 in the soma**

(A) Immunostaining analysis in FUS$^{WT}$ and FUS$^{P525L}$ hiPSC-derived spinal MNs growth cones (individual panels related to Figure 7D). GAP43 signal is magenta; PHALLOIDIN signal (marking growth cones) is green. Images on the right show the merge of GAP43, PHALLOIDIN and TYR-TUBULIN (tyrosinated alpha-tubulin; marking axons) (white). Scale bar: 10 μm. (B)

Immunostaining analysis in $FUS^{WT}$, $FUS^{P525L}$ and $FUS^{WT}$ overexpressing HuD under the SYN1 promoter ($FUS^{WT}$+HuD) hiPSC-derived spinal MNs. GAP43 signal is magenta; MAP2 signal is green. DAPI (blue) was used for nuclear staining. The graph shows the NRN1 signal intensity from 3 independent differentiation experiments, error bars indicate the , error bars indicate the standard error of the mean (Student's t test; unpaired; two tails; ****$p < 0.0001$). Scale bar: 10 μm.

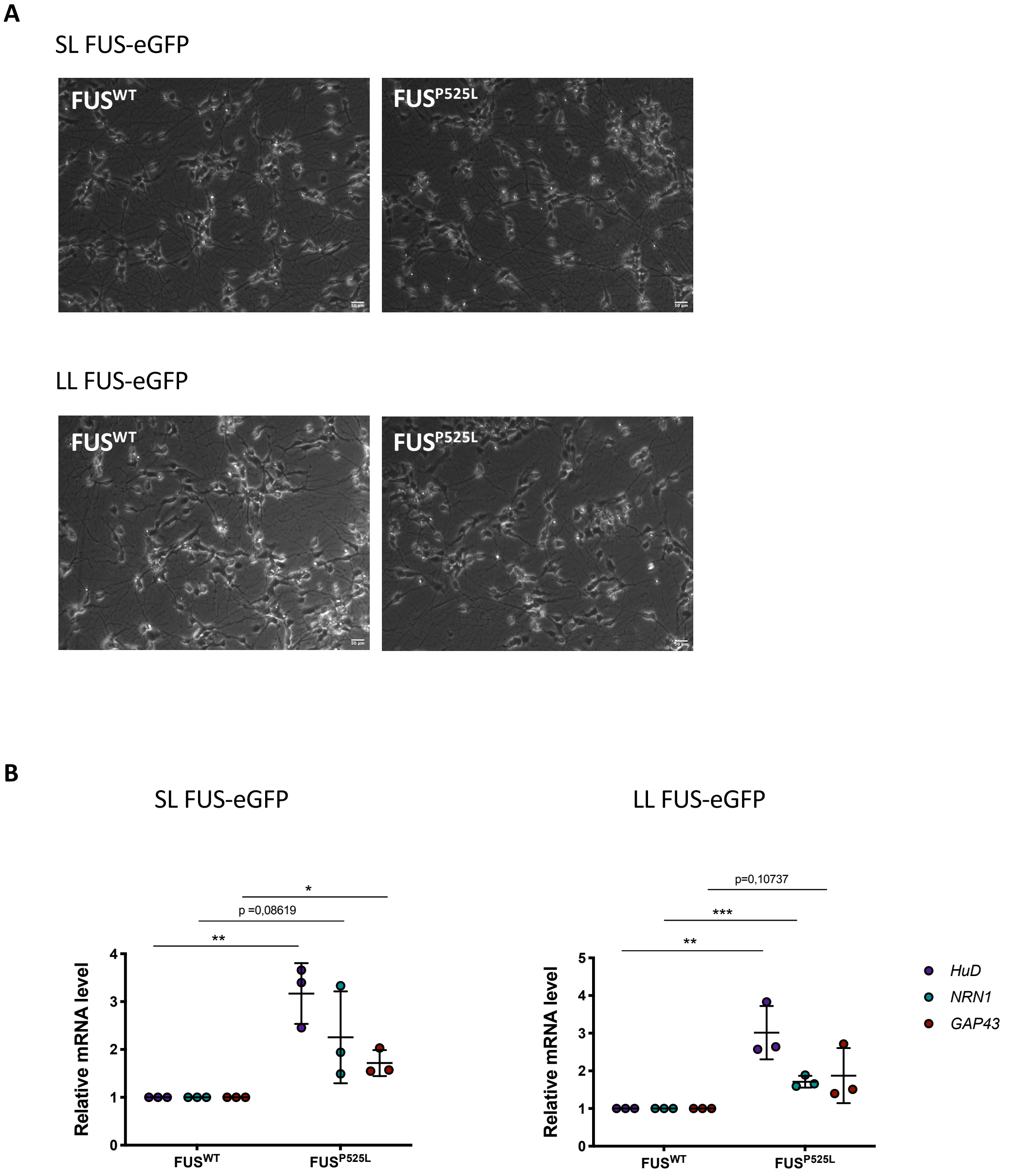


**Supplementary Figure S8. HuD, NRN1 and GAP43 mRNA levels in MNs from independent FUS mutant hiPSC lines**

(A) Phase contrast images of MNs obtained by differentiation of KOLF iPSCs WT 2 and P525L16 (LL FUS-eGFP) and T12.9 iPSCs WT15 and P525L17 (SL FUS-eGFP), a kind gift of J. Sterneckert [45]. Scale bars: 50 µm. (B) Analysis of the mRNA levels of the indicated genes by real time qRT-PCR in MNs shown in panel (A). The graph shows the average from 3 independent differentiation experiments, error bars indicate the standard deviation (Student's t-test; paired; two tails; *$p < 0.05$; **$p < 0.01$; ***$p < 0.001$).

**A**

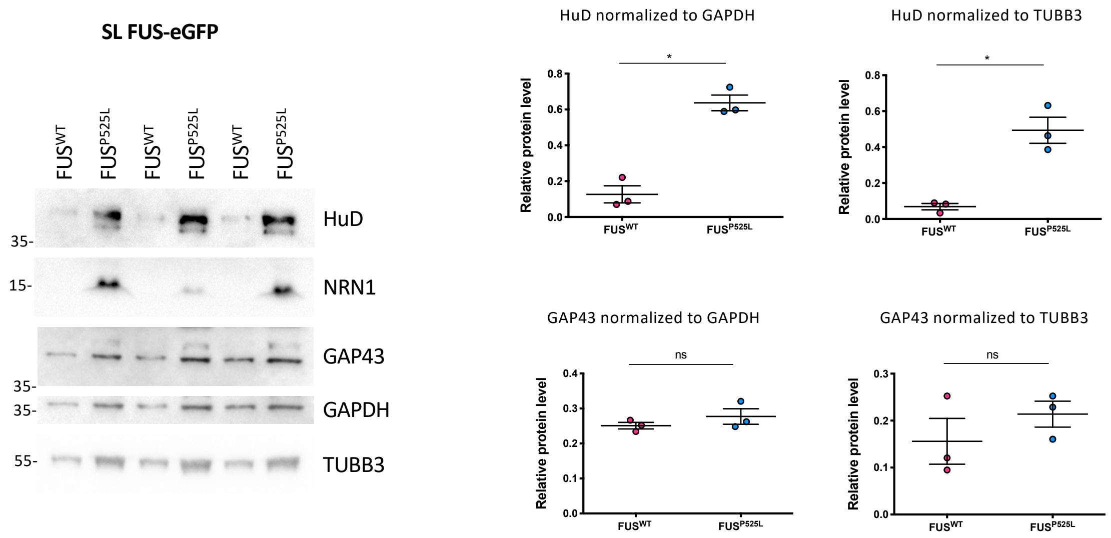


**B**

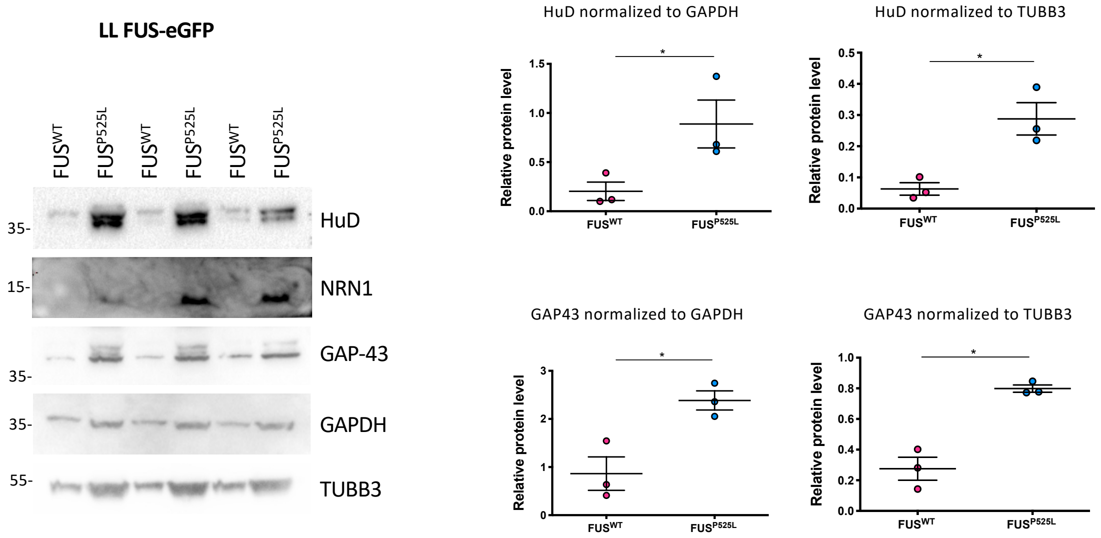


**Supplementary Figure S9. HuD, NRN1 and GAP43 protein levels in MNs from independent FUS mutant hiPSC lines**

(A,B) Analysis by western blot, of the levels of the indicated proteins in hiPSC-derived MNs shown in Supplementary Figure S6. Blots relating to 3 independent differentiation experiments are shown. The molecular weight is indicated on the left. The graphs show the quantification of the western blot signals normalized to TUBB3 or GAPDH, as indicated (averages and standard deviations; Student's t-test; paired; two tails; *$p < 0.05$; ns: $p > 0.05$).

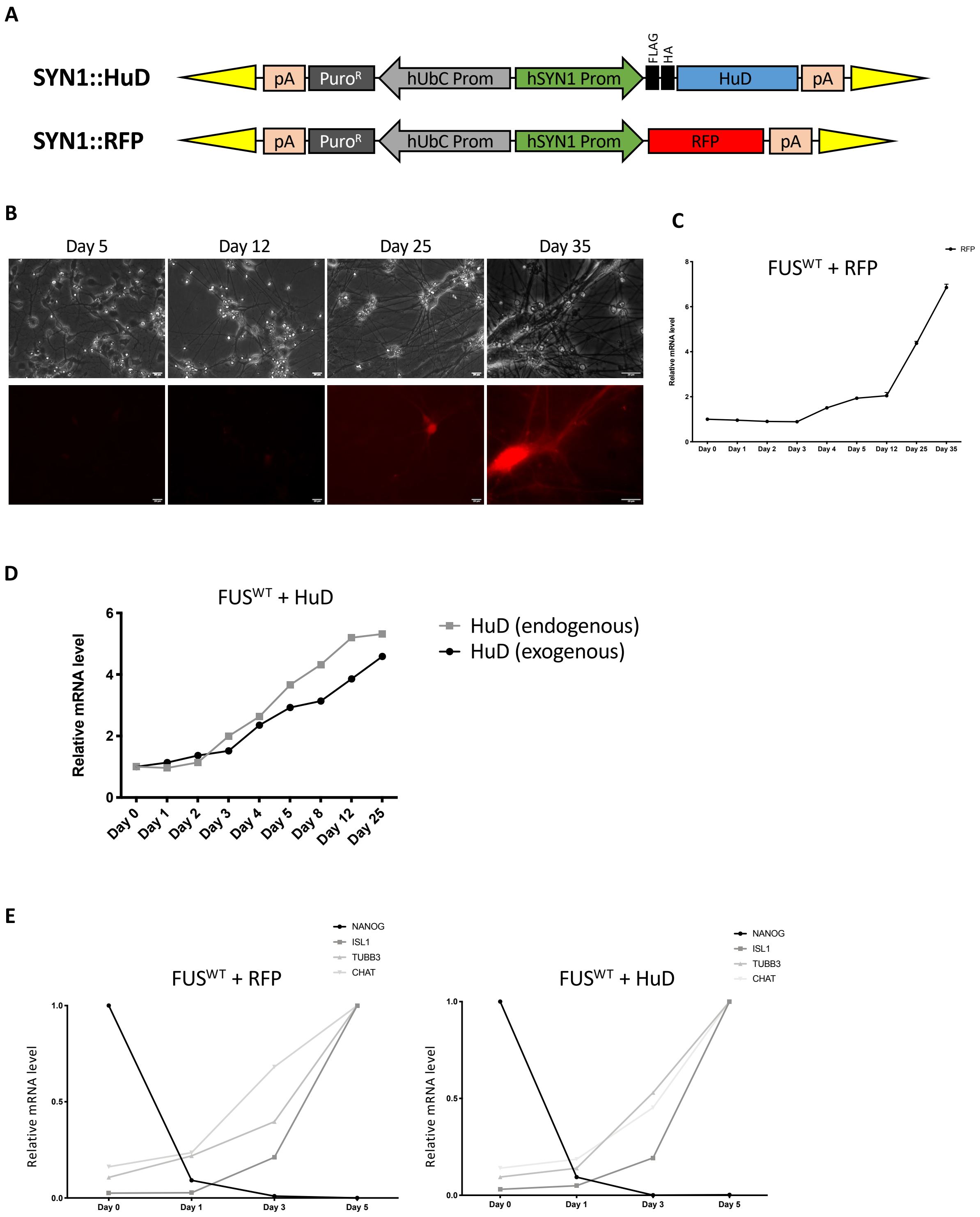


**Supplementary Figure S10. Validation of the SYN1::HuD construct**

(A) Schematic representation of the enhanced piggyBac (epB)-based vectors for the expression of HuD (top) or tagRFP (bottom) under the human synapsin 1 promoter (hSYN1 prom). Yellow triangles represent epB terminal repeats. pA: cleavage and polyadenylation site; $Puro^R$: puromycin resistance gene; hUbC Prom: constitutive promoter of the human Ubiquitin C gene. FLAG: DYKDDDDK tag. HA: human influenza hemagglutinin molecule tag. (B) Phase contrast (top) and RFP fluorescence (bottom) images of hiPSCs stably transduced with the tagRFP-encoding epB vector shown in panel A and induced to differentiate to MNs. Time points of differentiation are shown on top of the panels. Scale bars: 20 μm. (C) Real time qRT-PCR analysis of tagRFP expression in differentiating cells as in (B). (D) Expression of exogenous and endogenous HuD mRNA detected by real time qRT-PCR with specific primers. (E) Real time qRT-PCR analysis of the indicated marker genes in differentiating hiPSCs, showing that exogenous HuD expression does not alter MN differentiation.

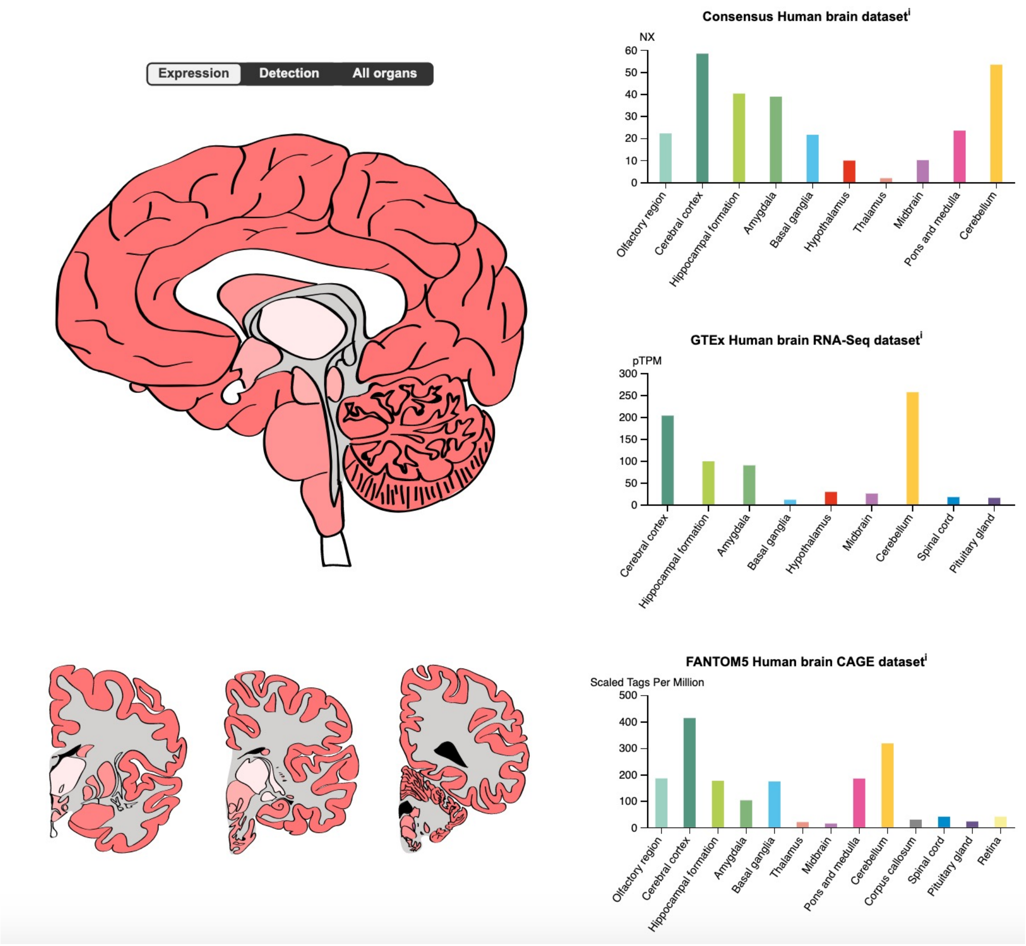


**Supplementary Figure S11. Expression of NRN1 in human nervous system**

Image and data available from v19.3.proteinatlas.org [65]. The Human Protein Atlas is available from http://www.proteinatlas.org and licensed under the Creative Commons Attribution-ShareAlike 3.0 International License.

| MGI | Symbol | Name | Annotated Term | Evidence | Inferred From | Reference(s) |
|---|---|---|---|---|---|---|
| 1099460 | RBM3 | RNA binding motif (RNP1, RRM) protein 3 | positive regulation of translation | ISO | RGD:620145 | J:155856 |
| 108177 | DHX9 | DEAH (Asp-Glu-Ala-His) box polypeptide 9 | positive regulation of cytoplasmic translation | ISO | Q08211 | J:164563 |
| 1100865 | RBM4 | RNA binding motif protein 4 | positive regulation of translation | IDA | | J:119742 |
| 1100851 | ELAVL1 | ELAV (embryonic lethal, abnormal vision)-like 1 (Hu antigen R) | positive regulation of translation | IDA | | J:175944 |
| 107427 | ELAVL4 | ELAV like RNA binding protein 4 | regulation of translation at synapse, modulating synaptic transmission | ISO | RGD:1560027 | J:155856 |
| 95564 | FMR1 | FMRP translational regulator 1 | negative regulation of cytoplasmic translation | IDA | | J:231207 |
| 107914 | TIA1 | cytotoxic granule-associated RNA binding protein 1 | negative regulation of translation | IMP | MGI:3037091 | J:88770 |

**Supplementary Table S2. Candidate regulators of HuD translation**

The table shows RBPs predicted to bind HuD 3’UTR by *cat*RAPID [35], which are also known regulators of translation. Gene Ontology Evidence Code Abbreviations: ISO, inferred from sequence orthology; IDA, inferred from direct assay; IMP, inferred from mutant phenotype.

| | |
|---|---|
| ATP5O FW | ACTCGGGTTTGACCTACAGC |
| ATP5O RV | GGTACTGAAGCATCGCACCT |
| HuD FW | CAACCCCAGCCAGAAGTCCA |
| HuD RV | AGCCTGAACCTCTGAGCCTG |
| NRN1 FW | GGCTTTTCGGACTGTTTGCTCA |
| NRN1 RV | ATCCTCCCAGTATGTGCACACG |
| SYN-Flag-HuD FW | ACTCAGCGCTGCCTCAGTCTG |
| SYN-Flag-HuD RV | ACCTGAGGCTCCATGGTGCTAAT |
| RFP FW | AACACTCGGCTGGGAGGCCAA |
| RFP RV | ACGTAGGTCTCTTTGTCGGCCT |
| NANOG FW | CCAAATTCTCCTGCCAGTGAC |
| NANOG RV | CACGTGGTTTCCAAACAAGAAA |
| TUJ1 FW | CCCGGAACCATGGACAGTGT |
| TUJ1 RV | TGACCCTTGGCCCAGTTGTT |
| CHAT FW | CTCAGCTACAAGGCCCTGCT |
| CHAT RV | ACCAGCGTGTCCTGGGTATG |
| ISL-1 FW | TACAAAGTTACCAGCCACC |
| ISL-1 RV | GGAAGTTGAGAGGACATTGA |
| GAP-43 FW | GAGGAGCCTAAACAAGCCGATG |
| GAP-43 RV | GGGCACTTTCCTTAGGTTTGGT |
| MAP1B FW | AGCCAGTCGAAGCCTACGTC |
| MAP1B RV | ATTCGCCCTCCCCTTCAGTG |
| F1_HuD-3'UTR-FW | TAATACGACTCACTATAGGGGCATTGAATGTTCTTTCATAGC |
| F1_HuD-3'UTR-RV | CTGTTTATAGCCCATCCTTC |
| F2_HuD-3'UTR-4-FW | TAATACGACTCACTATAGGGTGGGCTATAAACAGATGATCTT |
| F2_HuD-3'UTR-5-RV | CACAACAAACATGACAATATATCA |
| F3_HuD-3'UTR-6-FW | TAATACGACTCACTATAGGGGTTTGTGCTTTGTACGGTTA |
| F3_HuD-3'UTR-7-RV | CAATACTTTCCCTTGGAAATCATC |

**Supplementary Table S3. Sequences of the primers used for PCR or real-time qRT-PCR (indicated 5’ to 3’)**
